# STiLE: Automated Tissue Microarray Dearraying for Spatial Transcriptomics

**DOI:** 10.64898/2026.03.17.712359

**Authors:** Harsh Sinha, Arun Das, Yu-Chiao Chiu, Shou-Jiang Gao, Yufei Huang

## Abstract

Tissue microarrays (TMAs) enable high-throughput spatial transcriptomic profiling of dozens of tissue cores on a single slide. However, existing dearraying methods operate on histological images and do not support the coordinate-based outputs of spatial transcriptomics platforms. Therefore, task of assigning cells to their respective cores (dearraying) remains a manual bottleneck. We present STiLE, a tool for automated TMA dearraying that operates solely on cell centroid coordinates. By eliminating dependence on image data, STiLE is robust to artifacts such as variable staining quality and uneven illumination. The algorithm combines connectivity-based component detection, density-based clustering (HDBSCAN), component-guided cluster merging, and optional grid-based peak detection. Validation on eleven public TMA samples (50–150 cores, three platforms) achieved ARI > 0.99, while systematic benchmarking on 396 synthetic datasets with realistic artifacts demonstrated consistently robust performance (mean ARI = 0.992). STiLE accepts standard formats (AnnData, CSV) and is platform-agnostic, supporting diverse platforms including Vizgen MERSCOPE, 10x Xenium, and NanoString CosMx. An interactive Streamlit interface enables parameter tuning, visual inspection, and region-based processing for large slides.

**Availability and Implementation:** https://pypi.org/project/stile; **source at** https://github.com/Huang-AI4Medicine-Lab/stile

**Contact:** yu

**Supplementary Information:** Supplementary data are available at Bioinformatics online

## Introduction

Imaging-based spatial transcriptomics (iST) platforms such as 10x Xenium (5), NanoString CosMx (4), and Vizgen MERSCOPE (2) enable simultaneous profiling of hundreds to thousands of RNA targets *in situ* at subcellular resolution while preserving tissue architecture.These technologies provide unprecedented insight into spatial gene expression programs, tumor microenvironments, and cellular interactions. However, their high per-slide cost and limited field of view make large-cohort studies expensive and logistically challenging.

To address this constraint, many studies adopt tissue microarray (TMA) experimental designs (6; 8), in which dozens to hundreds of tissue cores from different patients are arrayed onto a single slide. When combined with iST, TMAs enable cohort-scale spatial profiling while amortizing imaging cost across many specimens (12). This strategy generates datasets containing millions of spatially resolved cells across large patient populations.

A critical preprocessing step in TMA-based iST is dearraying: assigning each segmented cell to its source core before downstream analysis. Accurate core assignment is essential for per-patient analyses, differential expression, spatial statistics, and integrative clinical modeling. Despite its importance, dearraying remains a practical bottleneck.

Existing dearraying approaches have been developed primarily for histological and multiplexed protein imaging workflows. Tools such as QuPath (1), ATMAD (9), Coreograph within MCMICRO (10), TMA-Grid (3), and PRISM (11) operate on raster images, detecting core boundaries through morphological features or deep learning–based segmentation. These methods assume dense, spatially continuous images with consistent tissue–background contrast, as produced by brightfield histology or multiplexed immunofluorescence platforms (e.g., CODEX, CyCIF, mIHC).

However, iST platforms differ fundamentally in both data structure and analytical workflow. Although fluorescence images are acquired during assay processing, standard downstream analysis relies on coordinate-based outputs consisting of cell centroids with associated transcript counts. Moreover, the fluorescence signals in iST are sparse and marker-dependent, lacking the uniform tissue–background contrast expected by existing image-based dearraying algorithms. In many experimental settings, matched histology images from the same slide may not be available, making image-driven TMA dearraying infeasible. Applying existing tools would require reverting to raw image data, reintroducing sensitivity to staining variability, uneven illumination, & tissue artifacts, despite the fact that coordinate data alone contains sufficient geometric information to delineate cores.

Furthermore, most image-based methods assume a regular grid layout, an assumption that breaks down when cores are misaligned, variably spaced, partially damaged, or missing. Grid drift and non-uniform spacing are common in large TMAs, particularly in slides subjected to sectioning artifacts or deformation. These limitations highlight a methodological gap: no existing dearraying framework is designed to operate directly on coordinate-based spatial transcriptomics outputs.

To address this need, we developed STiLE (Spatial Tissue microarray Labeling and Extraction), a Python tool that performs automated TMA dearraying directly from cell centroid coordinates. By operating entirely in coordinate space, STiLE eliminates dependence on histological images and is intrinsically robust to staining variability, illumination artifacts, and platform-specific imaging characteristics. Rather than imposing a rigid grid model, STiLE leverages connectivity-based component detection and density-based clustering to identify spatially coherent cores based on intrinsic geometric structure. An optional grid-aware refinement step regularizes core positions when grid structure is present, while preserving flexibility under irregular layouts.

STiLE is platform-agnostic, integrates directly with AnnData-based workflows, and scales to million-cell datasets. To our knowledge, it is the first dearraying method designed specifically for coordinate-based spatial transcriptomics data. By removing a persistent preprocessing bottleneck, STiLE enables scalable, cohort-level spatial transcriptomic analysis using cost-efficient TMA experimental designs.

### Implementation

STiLE operates through a modular, multi-phase pipeline designed to robustly separate spatially coherent TMA cores using only cell centroid coordinates. It consists of four stages: connectivity analysis, density-based clustering, and optional grid-based refinement (Fig. 1).

**Figure 1.**
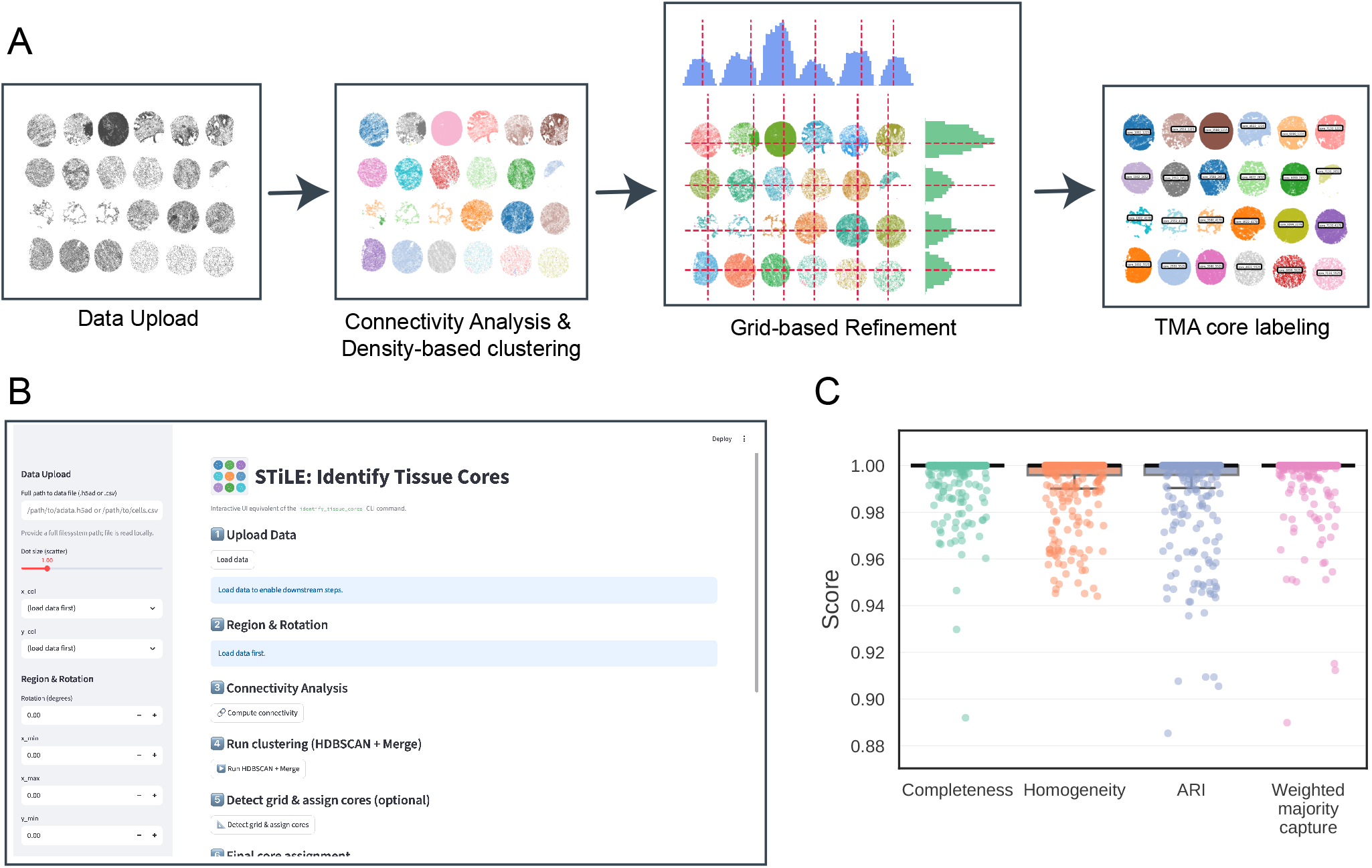
Overview of the STiLE dearraying workflow, interactive interface, and robustness evaluation. **(A)** Algorithmic pipeline. Cell centroids from spatial transcriptomics TMA data (CSV or AnnData) are first grouped into spatially coherent clusters via connectivity analysis on buffered geometric unions, followed by density-based clustering (HDBSCAN). An optional grid-based refinement step detects peaks in marginal density histograms to regularize core positions, after which cells are reassigned to their nearest core center. For large slides, the pipeline can be applied to sub-regions independently and merged. **(B)** Streamlit-based interactive interface for parameter adjustment, real-time visualization, validation, and sub-region processing. **(C)** Systematic robustness assessment across 396 synthetic TMA datasets incorporating realistic artifacts (missing cores, variable core sizes etc.). Each point represents one synthetic dataset across several metrics, completeness, homogeneity, adjusted Rand Index & weighted majority capture.

#### Connectivity analysis

The first phase identifies spatially separated coarse regions that correspond to candidate TMA cores. Each cell centroid is expanded by a buffer radius *r*, and cells whose buffered regions overlap are connected to form an undirected graph. Two circular buffers centered at points *p*_*i*_ and *p*_*j*_ overlap if and only if |*p*_*i*_ − *p*_*j*_ | ≤ 2*r*. Edges are computed efficiently using KD-tree–based neighbor queries, returning all pairs (*i, j*) satisfying the distance constraint in *O*(*n* log *n* + | 𝒫|) time, where | 𝒫| is the number of proximal pairs. The resulting sparse adjacency matrix is traversed using breadth-first search to extract connected components.

The default buffer radius *r* is defined as the median nearestneighbor distance estimated from a random subsample of up to 50,000 cells. This statistic provides a robust estimate of the intrinsic intra-core spatial scale. Within a TMA core, cells are densely packed, so the median nearest-neighbor distance captures typical intra-core spacing. In contrast, inter-core gaps are typically substantially larger than 2*r*, ensuring that cells from distinct cores fall into separate connected components. Users may override the default radius in rare cases where inter-core gaps approach intra-core spacing (e.g., nearly touching cores).

#### Density-based clustering

Connectivity analysis separates spatially disconnected regions but may retain sparse debris or segmentation artifacts within components. To refine core structure, STiLE applies HDBSCAN (Hierarchical Density-Based Spatial Clustering of Applications with Noise) with leaf cluster selection (7) independently within each connected component. Leaf-cluster selection favors compact, high-density clusters and reduces the risk of merging adjacent cores connected through sparse bridges. Noise points, including debris and low-density inter-core cells, are labeled as outliers and excluded from initial core assignments. Because tissue folding, partial sectioning, or density gradients may cause HDBSCAN to partition a single biological core into multiple subclusters, final consolidation is deferred to the next phase.

#### Component-guided merging

HDBSCAN may partition a single core into multiple sub-clusters due to density variation. STiLE resolves this by merging all clusters that share a connected component from the first phase. This component-guided merging restores biologically coherent cores while preserving the noise filtering benefits of density-based clustering. For well-separated TMAs, this step typically yields final core assignments without further processing.

#### Grid-based refinement (optional)

In challenging cases, such as sparse cores, fragmented tissue, or narrow inter-core gaps, connectivity analysis alone may yield fragmented or partially merged cores. To address this, STiLE provides an optional grid-aware refinement step that leverages global array structure without imposing a rigid grid assumption. Specifically, cell coordinates are projected onto marginal density histograms along the *x* and *y* axes (Fig. 1A). Peaks are detected using prominence-based criteria, ranked by the product of peak prominence and histogram height. The top *k*_*x*_ and *k*_*y*_ peaks define candidate grid coordinates along each axis. The cartesian product of these peak sets yields candidate core centers. Merged clusters are then reassigned to their nearest candidate center. Candidate centers with insufficient support (below a minimum cell count threshold) are discarded. This hybrid approach preserves the adaptability of connectivity clustering while leveraging the grid structure when available.

#### Interactive interface

For large slides with heterogeneous spacing, users may define rectangular sub-regions, process each independently, and merge results with region-specific labels. STiLE provides a Streamlit-based web application that can be deployed locally or on institutional servers, avoiding external data transfer. It supports the complete workflow, including parameter selection, data upload, region selection, parameter adjustment for each phase, real-time visualization, and export of annotated results.

## Results

We validated STiLE on publicly available TMA datasets from (12), comprising tumor and normal tissue arrays profiled on three imaging-based spatial transcriptomics platforms: 10x Xenium, NanoString CosMx, and Vizgen MERSCOPE (Table 1). Ground truth core identities were derived from the original TMA layout maps. Because STiLE assigns arbitrary core indices that do not correspond to ground truth labels, we used the Hungarian algorithm to find the optimal one-to-one matching between predicted and true core assignments before computing evaluation metrics.

**Table 1.** Validation on public TMA datasets (12). ARI: Adjusted Rand Index, Comp: Completeness, Hg: Homogeneity, WMC: Weighted majority capture.

| Tissue | TMA | Platform | Cores | Cells | Comp. | Hg | ARI | WMC |
| --- | --- | --- | --- | --- | --- | --- | --- | --- |
| Multi-Tissue | Normal | CosMx | 47 | 178,424 | 0.99 | 0.99 | 0.99 | 0.99 |
| Multi-Tissue | Tumor | CosMx | 163 | 240,154 | 1.00 | 1.00 | 1.00 | 1.00 |
| Lung | Tumor | Xenium | 149 | 360,376 | 1.00 | 1.00 | 1.00 | 1.00 |
| Pan-human | Normal | Xenium | 48 | 290,234 | 0.99 | 0.99 | 0.99 | 0.99 |
| Pan-human | Tumor | Xenium | 159 | 379,365 | 0.99 | 1.00 | 0.99 | 0.99 |
| Multitissue | Tumor TMA 1 | CosMx | 144 | 258,986 | 0.99 | 1.00 | 0.99 | 0.99 |
| Multitissue | Tumor TMA 2 | CosMx | 47 | 249,682 | 1.00 | 1.00 | 1.00 | 1.00 |
| Breast | Tumor TMA 1 | MERSCOPE | 89 | 154,154 | 0.99 | 1.00 | 0.99 | 0.99 |
| Breast | Tumor TMA 2 | MERSCOPE | 44 | 125,573 | 0.99 | 0.99 | 0.99 | 0.99 |
| Breast | Tumor TMA 1 | Xenium | 145 | 342,704 | 1.00 | 1.00 | 1.00 | 1.00 |
| Breast | Tumor TMA 2 | Xenium | 49 | 299,900 | 0.99 | 0.99 | 0.99 | 0.99 |

STiLE achieved high accuracy across all datasets, with completeness, homogeneity, Adjusted Rand Index (ARI), and weighted majority capture exceeding 0.99 in each case (Table 1). Processing time scaled approximately as *O*(*n* log *n*) with cell count *n*, dominated by HDBSCAN clustering and kd-tree construction; datasets with up to one million cells were completed within a couple of minutes on standard hardware.

To systematically assess robustness under controlled conditions, we generated 396 synthetic TMA datasets with realistic artifacts. The simulations varied with three primary parameters: core size irregularity (radius jitter 0–100%), cell density bias across cores (0–100%), and core missingness (0–50%). All synthetic datasets additionally included elliptical core deformation, tissue artifacts (slits, bubbles, folds), affine warping, and non-rigid thin-plate-spline deformation to mimic real-world variability. Across all 396 configurations, STiLE achieved a mean ARI of 0.992 (median 1.000, minimum 0.885), with 99.7% of datasets exceeding ARI 0.90 (Supplementary Tables 6–8). Performance remained stable under challenging conditions: mean ARI was 0.991 even at 100% radius jitter, and degradation was minimal with up to 50% missing cores. These results demonstrate that STiLE’s connectivity-based approach is robust to the geometric and density variations commonly encountered in real TMA preparations.

## Discussion

STiLE resolves a persistent preprocessing bottleneck in TMA-based spatial transcriptomics by enabling automated dearraying directly from cell centroid coordinates. By operating entirely in coordinate space, it avoids dependence on histological image quality, staining variability, and platform-specific imaging, unlike imagebased methods such as MCMICRO (10) and ATMAD (9). This makes STiLE immediately applicable to platforms including 10x Xenium, NanoString CosMx, and Vizgen MERSCOPE without retraining or image preprocessing.

Its modular design adapts to diverse TMA configurations. For well-separated cores, connectivity analysis with density-based clustering yields accurate assignments. For dense or irregular arrays, optional grid-based refinement introduces global structural guidance without enforcing a rigid grid. Region-based processing further supports large slides with heterogeneous spacing, allowing flexibility while maintaining robustness.Unlike end-to-end systems such as PRISM (11), STiLE focuses on dearraying as a modular component, producing interoperable outputs that integrate into existing workflows while eliminating a manual preprocessing step.

Limitations include the assumption that inter-core gaps exhibit lower cell density than intra-core regions; touching or extremely sparse cores may require manual adjustment. STiLE also assumes upstream cell segmentation has been completed and is therefore complementary to broader analysis frameworks.

Overall, by reframing TMA dearraying as a geometric clustering problem in coordinate space, STiLE provides a scalable and platform-agnostic solution for cohort-scale spatial transcriptomic analysis.

## Funding

We thank members of Drs. Shou-Jiang Gao and Yufei Huang laboratories for technical assistance and discussions. This study was supported by grants from the National Institutes of Health (CA096512, CA284554, CA278812, CA291244 and CA124332 to S.-J. Gao; U01CA279618 and R21GM155774 to Y. Huang; R35GM154967 to Y.-C. Chiu), UPMC Hillman Cancer Center Startup Funds to S.-J. Gao and Y. Huang, and in part by award P30CA047904. This research was also supported in part by the University of Pittsburgh Center for Research Computing, RRID:SCR 022735. Specifically, this work used the HTC cluster, which is supported by S10OD028483.

## Data availability

No new experimental data were generated for this study. Public TMA datasets used for validation were obtained from (12)[10.5281/zenodo.16847475]. Simulated TMA datasets and STile-generated core annotations for all public datasets are available at Zenodo [10.5281/zenodo.18880122]

## Author contributions statement

Conceptualization, H.S., A.D., Y.H.; Data curation, H.S., A.D.; Investigation, Methodology, Software, Validation, Visualization, H.S.; Writing-original draft, H.S., A.D., Y.H.; Writing - review & editing, H.S., A.D., Y.-C.C., S.-J.G., Y.H.; Funding acquisition, Y.-C.C., S.-J.G., Y.H.; Supervision, Y.H.

## Supplementary Materials

### S1 Algorithm Details

#### S1.1 Pipeline Pseudocode

Algorithm 1 provides a formal description of the STiLE pipeline. The five phases are applied sequentially, with Phases 4–5 being optional depending on data characteristics.

##### Algorithm 1

STiLE Pipeline

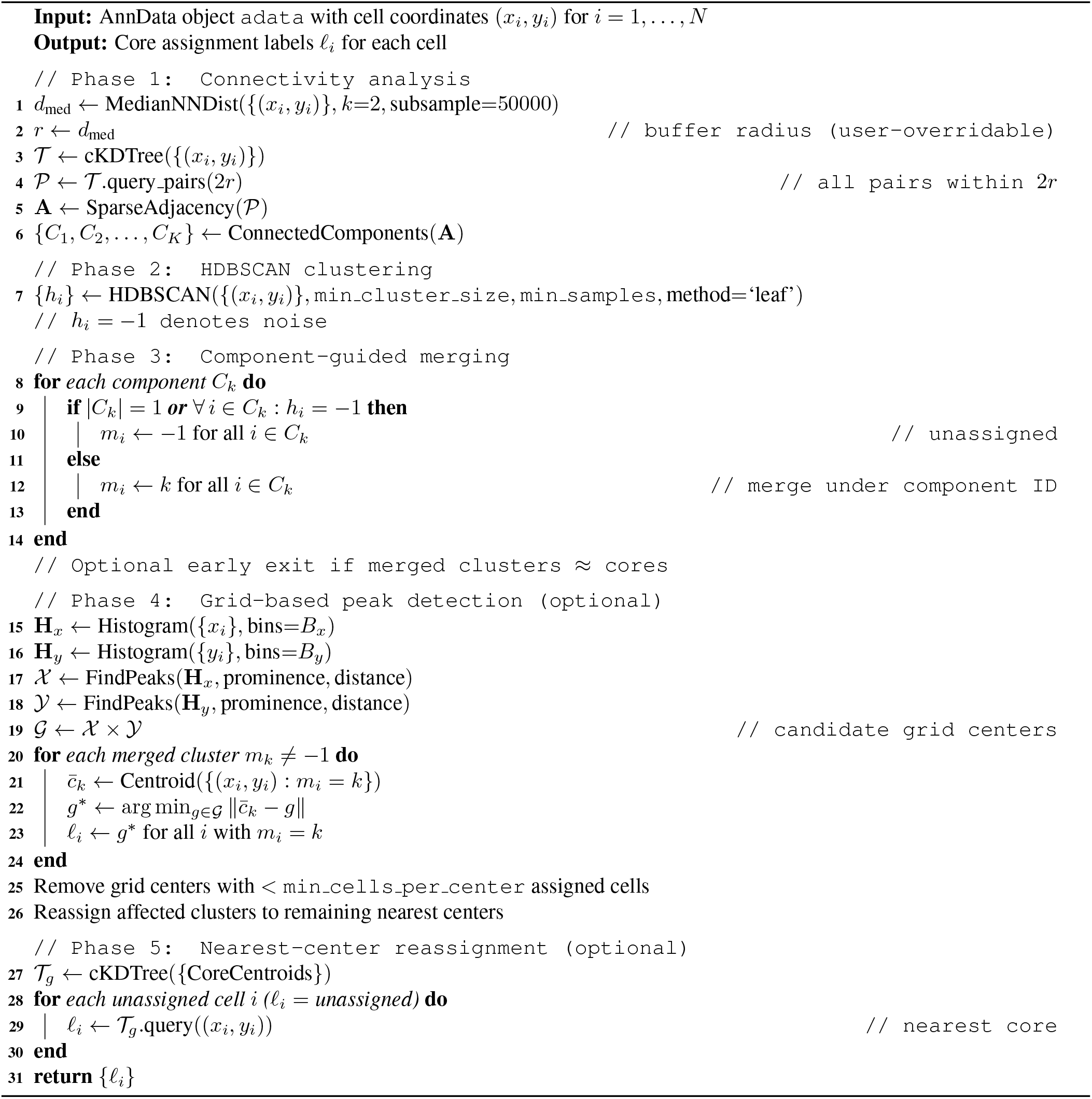

### S2 Parameter Reference

Table 1 lists all user-adjustable parameters, their defaults in both the CLI and interactive Streamlit interface, and their effects on pipeline behavior.

**Table 1.**
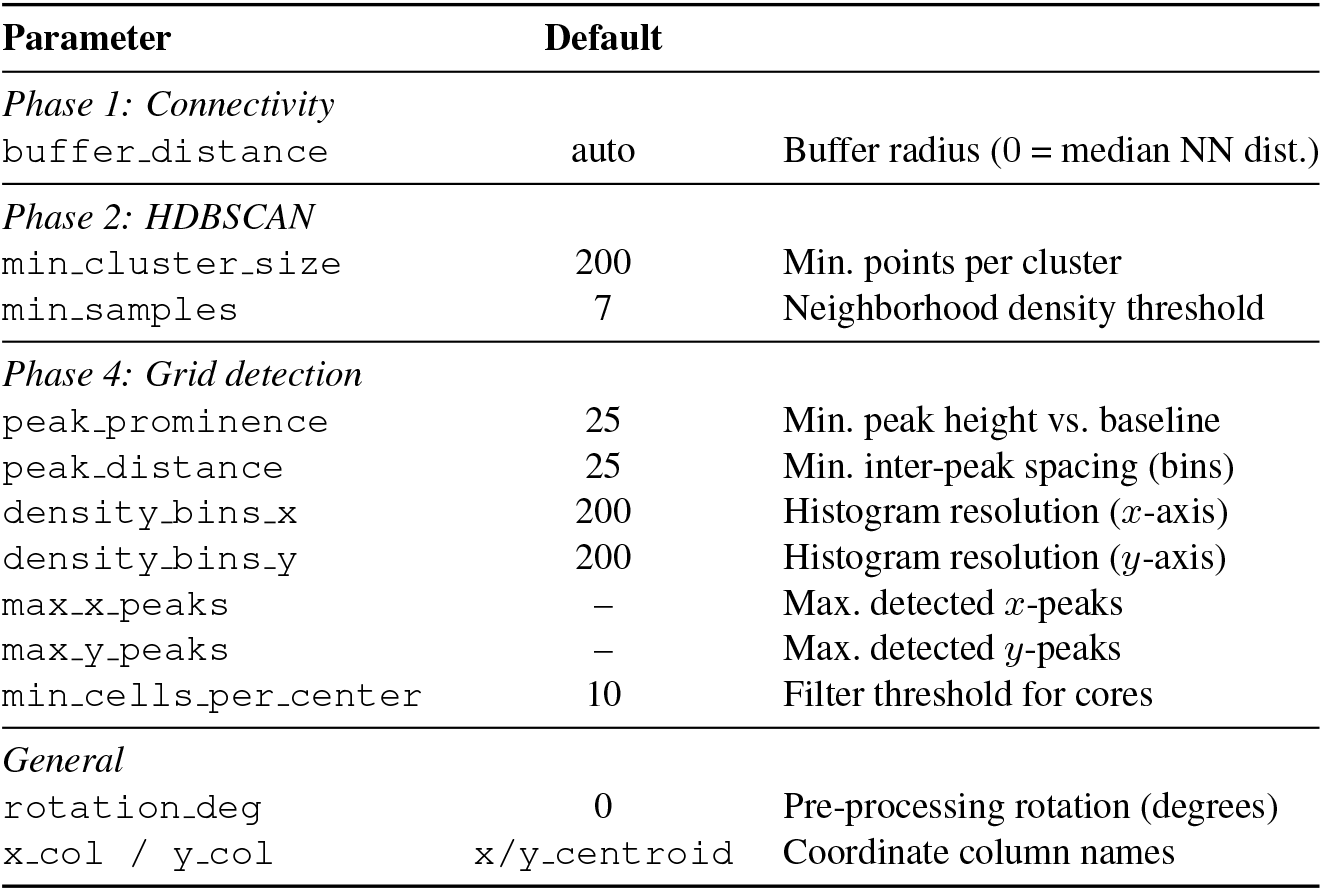
Complete parameter reference.

#### Parameter guidance

For well-separated TMAs with 20–50 cores, the defaults generally suffice. For dense TMAs with *>*100 cores, we recommend: (i) reducing `peak_prominence` and `peak_distance` to detect closely spaced peaks, (ii) using the region-based workflow to limit peak detection to sub-regions with locally uniform spacing, and (iii) increasing `min_cluster_size` proportionally to the cell density to avoid over-fragmentation.

#### S2.1 Processing Time

Table 2 reports wall-clock processing times on a standard workstation (Intel Xeon, 16 GB RAM, single-threaded except for HDBSCAN core distance computation with 4 threads).

**Table 2.** Processing time by dataset.

| TMA | Platform | Cells | Time (s) | Phase Breakdown |
| --- | --- | --- | --- | --- |
| Tumor | Xenium | 1.2M | 92 | P1: 18s, P2: 45s, P3: 2s, P4: 15s, P5: 12s |
| Tumor | CosMx | 0.8M | 68 | P1: 12s, P2: 33s, P3: 1s, P4: 12s, P5: 10s |
| Normal | Xenium | 0.4M | 24 | P1: 5s, P2: 14s, P3: 1s |
| Normal | MERSCOPE | 0.3M | 18 | P1: 4s, P2: 10s, P3: 1s |

HDBSCAN clustering (Phase 2) dominates the processing time, scaling approximately as *O*(*n* log *n*) in the number of cells. The connectivity analysis (Phase 1) is the second-most expensive step due to the kd-tree pair query. Grid detection (Phase 4) and reassignment (Phase 5) are comparatively fast.

#### S2.2 Simulation Benchmark

We systematically evaluated STiLE on 396 synthetic TMA datasets generated with the built-in simulation module (`stile.simulation`). Datasets varied three difficulty axes—radius jitter fraction (0–100%, 11 levels), radial density bias (0–100%, 6 levels), and missing core fraction (0–50%, 6 levels)—in a full factorial design. All datasets included elliptical cores, tissue slits (60% of cores), bubbles (8), folds (3), affine warping (rotation + scale + shear), thin-plate-spline non-rigid deformation (amplitude 100), edge cropping, and block/random missingness. Tables 3–5 report ARI stratified by each axis.

**Table 3.** ARI by radius jitter fraction. Each level contains 36 datasets.

| Jitter (%) | $n$ | ARI mean | ARI min |
| --- | --- | --- | --- |
| 0 | 36 | 0.9946 | 0.9356 |
| 10 | 36 | 0.9987 | 0.9904 |
| 20 | 36 | 0.9988 | 0.9917 |
| 30 | 36 | 0.9925 | 0.9094 |
| 40 | 36 | 0.9928 | 0.9094 |
| 50 | 36 | 0.9880 | 0.9494 |
| 60 | 36 | 0.9833 | 0.8853 |
| 70 | 36 | 0.9919 | 0.9468 |
| 80 | 36 | 0.9901 | 0.9076 |
| 90 | 36 | 0.9921 | 0.9451 |
| 100 | 36 | 0.9905 | 0.9416 |

Across all 396 datasets, STiLE achieved a mean ARI of 0.992 (median 1.000, minimum 0.885), with 395 of 396 datasets (99.7%) exceeding ARI 0.90. Performance remained consistently high across all jitter levels (mean ARI ≥ 0.983), density bias levels (mean ARI ≥ 0.987), and missingness levels (mean ARI ≥ 0.970).

**Table 4.** ARI by radial density bias. Each level contains 66 datasets.

| Bias (%) | $n$ | ARI mean | ARI min |
| --- | --- | --- | --- |
| 0 | 66 | 0.9925 | 0.9356 |
| 20 | 66 | 0.9919 | 0.9434 |
| 40 | 66 | 0.9937 | 0.9447 |
| 60 | 66 | 0.9941 | 0.9416 |
| 80 | 66 | 0.9936 | 0.9471 |
| 100 | 66 | 0.9869 | 0.8853 |

**Table 5.** ARI by missing core fraction. Each level contains 66 datasets.

| Missing (%) | $n$ | ARI mean | ARI min |
| --- | --- | --- | --- |
| 0 | 66 | 0.9705 | 0.9369 |
| 10 | 66 | 0.9902 | 0.9055 |
| 20 | 66 | 1.0000 | 0.9995 |
| 30 | 66 | 1.0000 | 1.0000 |
| 40 | 66 | 0.9921 | 0.8853 |
| 50 | 66 | 1.0000 | 0.9992 |

### S3 Region-Based Workflow

For large TMA slides with many cores (*>*50), global peak detection on marginal density histograms can fail because:

1. **Non-uniform spacing**: Core pitch may vary across the slide due to manufacturing tolerances, causing peaks at inconsistent intervals.
2. **Peak merging**: Adjacent cores may produce overlapping peaks in the marginal histogram, leading to under-detection.
3. **Missing cores**: Absent cores create gaps in the expected grid, breaking the assumption of uniform peak spacing.

STiLE’s region-based workflow addresses these challenges:

1. **Region definition**: The user defines a rectangular bounding box (*x*_min_, *x*_max_, *y*_min_, *y*_max_) covering a subset of the TMA.
2. **Cropping**: Cells outside the bounding box are excluded. The cropped dataset retains original cell indices for later merging.
3. **Independent processing**: The full pipeline (Phases 1–5) is applied to the cropped region. Within each region, core spacing is locally consistent, enabling reliable peak detection.
4. **Region labeling**: Core labels are prefixed with the region name (e.g., `region1_core_1234_5678`) to prevent collisions across regions.
5. **Persistence**: Region results are saved as CSV files in a dataset-specific directory, enabling incremental processing across sessions.
6. **Merging**: After all regions are processed, a merge step combines results by matching cell indices and writing each region’s prefixed core labels back to the original dataset.

**Figure 1.**
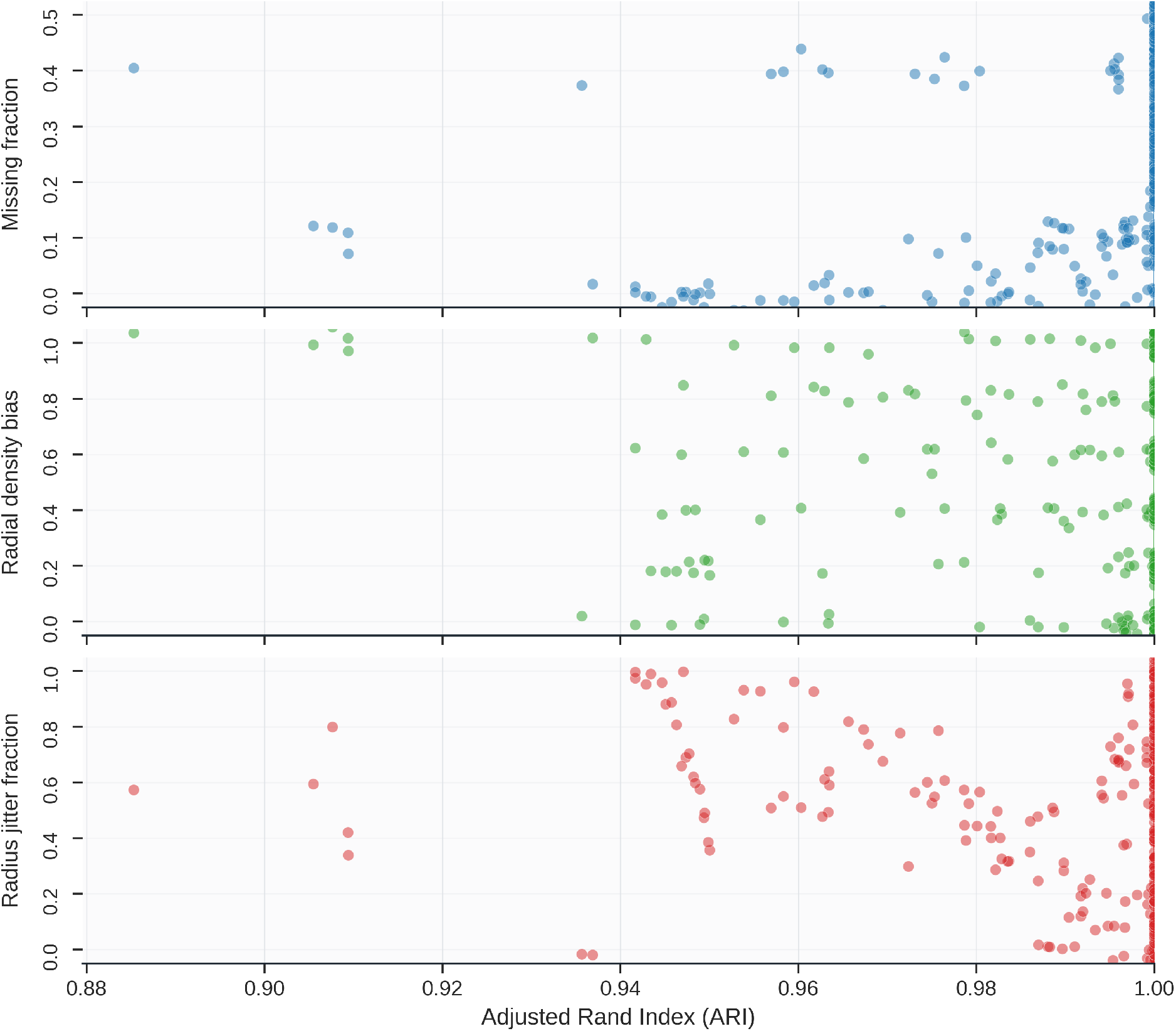
Evaluation on simulated TMA datasets designed to mimic realistic TMA datasets. Each simulated dataset contains elliptical cores with varying levels of controlled structural perturbations, including missing cores, radial density bias and radius jitter. Each point represents a single simulated dataset and the performance is measured using the Adjusted Rand Index (ARI)

The Streamlit interface provides visual guidance for region selection, displaying the current cell scatter plot with adjustable bounding box sliders.

### S4 Interactive Interface

The Streamlit-based interface (`stile run`) provides a nine-step guided workflow:

1. **Data upload**: Load `.h5ad` or `.csv` files. For CSV files, users select the *x* and *y* coordinate columns. Optional rotation applies an affine transformation before processing.
2. **Region cropping**: Define rectangular bounds via sliders to restrict analysis to a sub-region. A scatter plot previews the selected region.
3. **Connectivity analysis**: Compute median nearest-neighbor distance and identify connected components. Displays the number of components found.
4. **HDBSCAN + merge**: Run HDBSCAN clustering and component-guided merging. Sidebar controls adjust `min_cluster_size` and `min_samples`. A scatter plot colored by merged cluster label provides immediate visual feedback.
5. **Grid detection**: Construct marginal density histograms and detect peaks. Interactive histograms with peak markers allow users to assess detection quality. Sidebar controls adjust prominence, distance, and bin count.
6. **Final assignment**: Review core assignments on a labeled scatter plot. Save the current region’s results (with optional region name prefix).
7. **Region merging**: Select and merge previously saved regions into the full dataset.
8. **Export**: Download core assignments as CSV or save the annotated `.h5ad` file.
9. **Layout mapping**: Upload a layout CSV defining the desired core naming scheme (e.g., A01, A02, …). STiLE maps detected core positions to the layout grid and creates a `core_name` column.

All parameters persist across workflow steps via Streamlit session state. The interface supports iterative refinement: users can return to any step, adjust parameters, and re-run downstream steps.

#### S4.1 Programmatic Usage

For scripting and integration into custom workflows, the pipeline can be invoked directly from Python:

`from stile.pipeline import identify_tissue_cores import scanpy as sc`

`adata = sc.read_h5ad(“data.h5ad”)`

`adata = identify_tissue_cores(adata)`

Individual phase functions (`find_connected_components`, `assign_cells_hdbscan`, `merge_clusters_by_component`, etc.) are importable from `stile.nodes` for fine-grained control.

### S5 Data Format Conventions

#### S5.1 Input Requirements

STiLE accepts data in two formats:

- **AnnData** (`.h5ad`): Cell coordinates must be present as numeric columns in `adata.obs`. Default column names are `x_centroid` and `y_centroid`, configurable via `x_col` and `y_col` parameters.
- **CSV**: Must contain at least two numeric columns representing spatial coordinates. The user selects the appropriate columns in the Streamlit interface.

Platform-specific raw data can be converted to standardized format using `stile prepare_data`:

- **Vizgen MERSCOPE**: Reads `cell_by_gene.csv` and `cell_metadata.csv`.
- **10x Xenium**: Reads `cell_feature_matrix.h5` and `cells.csv`.
- **NanoString CosMx**: Reads `exprMat_file.csv` and `cell_metadata.csv`.

#### S5.2 Output Format

Core assignments are stored in `adata.obs` with the following columns:

- `hdbscan_cluster` (int): HDBSCAN cluster label ( − 1 = noise).
- `connected_component` (int): Connected component ID.
- `merged_cluster` (int): Component-merged cluster label ( − 1 = unassigned).
- `path_block_core` (str): Final core label, formatted as `core_{x}_ {y}` where *x* and *y* are the integer coordinates of the assigned core center, or `unassigned`.
- `path_block_core_x`, `path_block_core_y` (float): Coordinates of the assigned core center.
- `core_name` (str, optional): Human-readable core name from layout mapping.

### S6 Evaluation Metrics

- **Adjusted Rand Index (ARI)**: Measures agreement between predicted and true clusterings, corrected for chance. Range [− 1, 1]; 1 indicates perfect agreement.
- **Homogeneity**: Each predicted cluster contains only members of a single true cluster.
- **Completeness**: All members of a true cluster are assigned to the same predicted cluster.
- **Weighted Majority Capture**: For each true cluster, the fraction of its cells assigned to the most common predicted cluster, weighted by cluster size. Values near 1 indicate low fragmentation.

### S7 Synthetic Data Generation

We used simulation for generating synthetic TMA datasets with configurable artifacts. This enables systematic benchmarking across controlled conditions.

#### Grid configuration

Users specify the number of rows and columns, inter-core pitch, core radius, and origin. Core positions are subject to center jitter, and core shapes can be elliptical with configurable eccentricity and orientation.

#### Cell sampling

Cells are sampled within each core from a distribution controlled by `MEAN_CELLS_PER_CORE` (Poisson-distributed) and `RADIAL_DENSITY_BIAS` (center-biased density).

#### Artifact simulation

- **Slits**: Linear cracks through cores, parameterized by probability, count, width, and angle.
- **Missing cores**: Random core dropout and block (row/column) dropout.
- **Bubbles**: Circular regions of cell removal, simulating air bubbles.
- **Folds**: Curved bands of cell removal, simulating tissue folds.
- **Affine deformations**: Global rotation, scaling, shear, and translation.
- **Thin-plate spline (TPS) warping**: Non-rigid deformation with configurable amplitude and control points.

The simulation outputs cell coordinates with ground-truth core assignments, enabling automated evaluation via the metrics module.

## Notes

### Competing Interest Statement

The authors have declared no competing interest.

